# Senescent cell heterogeneity and responses to senolytic treatment are related to cell cycle status during cell growth arrest

**DOI:** 10.1101/2024.06.22.600200

**Authors:** Francesco Neri, Shuyuan Zheng, Mark Watson, Pierre-Yves Desprez, Akos A. Gerencser, Judith Campisi, Denis Wirtz, Pei-Hsun Wu, Birgit Schilling

**Affiliations:** Buck Institute for Research on Aging, Novato, CA, USA; USC Leonard Davis School of Gerontology, Los Angeles, CA, USA; Department of Chemical and Biomolecular Engineering, Johns Hopkins University, Baltimore, MD, USA; California Pacific Medical Center, San Francisco, CA, USA; Johns Hopkins Physical Sciences–Oncology Center and Institute for NanoBioTechnology, Johns Hopkins University, Baltimore, MD, USA

**Keywords:** Cellular senescence, high-content image analysis, DNA content, heterogeneity, senolytics, cell cycle

## Abstract

Cellular senescence has been strongly linked to aging and age-related diseases. It is well established that the phenotype of senescent cells is highly heterogeneous and influenced by their cell type and senescence-inducing stimulus. Recent single-cell RNA-sequencing studies identified heterogeneity within senescent cell populations. However, proof of functional differences between such subpopulations is lacking. To identify functionally distinct senescent cell subpopulations, we employed high-content image analysis to measure senescence marker expression in primary human endothelial cells and fibroblasts. We found that G2-arrested senescent cells feature higher senescence marker expression than G1-arrested senescent cells. To investigate functional differences, we compared IL-6 secretion and response to ABT263 senolytic treatment in G1 and G2 senescent cells. We determined that G2-arrested senescent cells secrete more IL-6 and are more sensitive to ABT263 than G1-arrested cells. We hypothesize that cell cycle dependent DNA content is a key contributor to the heterogeneity within senescent cell populations. This study demonstrates the existence of functionally distinct senescent subpopulations even in culture. This data provides the first evidence of selective cell response to senolytic treatment among senescent cell subpopulations. Overall, this study emphasizes the importance of considering the senescent cell heterogeneity in the development of future senolytic therapies.

## Introduction

Cellular senescence is a complex stress response, generally characterized by an essentially irreversible cell cycle arrest, altered morphology, increased lysosomal activity, macromolecular damage, and profound changes in gene expression, such as the acquisition of a senescence-associated secretory phenotype (SASP)^1^. Genetic and pharmacological interventions that promote the elimination of senescent cells, a process termed senolysis, have shown to benefit the healthspan and lifespan of mice^2–4^. Because of their promising results in animal models, senolytic therapies have now entered clinical trials^5,6^. Notably, a senolytic drug tested in human patients with diabetic macular degeneration, UBX1325^7^, was recently shown to improve vision for up to 48 weeks after a single eye injection (local administration in a Phase 2 clinical trial^8^). While current treatments hold premise to improve selected age-related pathologies, much more work is needed to develop safe and effective senolytics that might have the potential to significantly improve human healthspan. In general, systemic treatments using senolytics might not be feasible as senescent cells also hold important functions such as wound healing. Thus, current studies often focus on local administration of senolytics in human clinical trials. Another important challenge however, in the development of senolytics (eliminating senescent cells) or senomorphics (suppressing the secretory phenotype, the SASP) is the heterogeneity of senescent cells, which is still poorly understood^9^.

Senescent cells are highly heterogeneous, as many different cell types in different organs can become senescent due to a variety of stressors capable of inducing senescence. Indeed, Coppe et al.^10^ and Basisty et al.^11^ showed that senescent cell culture models based on diverse cell types and senescence inducers resulted in senescent cells with different SASP. These different SASP profiles are influenced by the distinct gene expression profiles across these heterogeneous senescent cell populations^12^. However, several previous studies involved the use of bulk analytical techniques, and therefore could not decipher the heterogeneity within a given senescent population. Recently, single-cell RNA-sequencing (scRNA-Seq) has been used to analyze the diversity within senescent cell populations in culture revealing high levels of heterogeneity in cellular senescence^13–15^. Despite the carefully controlled uniform experimental conditions, significant heterogeneity was observed in different senescent cell clusters with different transcriptomic profiles, suggesting the existence of senescent cell subpopulations with different cell states that likely would exhibit distinct function and biology. Further research is therefore needed to characterize the actual functional differences between senescent cell subpopulations in culture, but eventually also in vivo on an organismal level.

Previous studies demonstrated that senolytics showed different efficacies across senescent cell culture models (based on different cell types and senescence inducers) using bulk RNA-Seq technologies ^4,16^. However, it remained unknown whether there is heterogeneity in senolytic responses within senescent cell populations. While this seems likely because of the existence of heterogeneous cell subpopulations highlighted by previous single-cell studies, no evidence of this is yet available. Understanding whether heterogeneous subpopulations of senescent cells indeed respond differently to senolytics is critical for the development of the next generation of senolytic (and/or senomorphic) therapies. Unlike scRNA-Seq, high-content imaging – which entails the automated acquisition and analysis of microscopic images from biological samples^17,18^ – enables rapid and cost-effective measurements of several senescence markers at a single-cell level. Even though imaging is limited by the number of markers assessed at once, it allows measurements at the protein level – one step closer to function than RNA. Additionally, imaging can readily assess cell viability and hence the response to senolytics. Thus, we set out to identify functionally distinct senescent cell subpopulations that might respond differently to senolytics using a high-content imaging workflow.

## Results

### Validation of senescence and heterogeneity of different cell populations

To study the heterogeneity of cellular senescence, the expression of several senescence-associated markers was measured at the single-cell level by using a high-content image analysis workflow (Figure 1). As a senescence model, primary human endothelial cells (HMVEC-L) and primary human fibroblasts (IMR-90) were cultured, and senescence was induced using ionizing radiation (IR) (Figure 1A). Both senescent (SEN) and their control samples (CTL) were either cultured in full-serum medium for the entire experiment (FS) or were switched to low-serum medium for the last 3 days of culture to induce quiescence by serum-starvation in CTL cells (SS). SEN and CTL samples were subsequently co-stained either for SA-β-Gal^19^ and proliferation (EdU incorporation) or for other senescence markers (γH2AX, LaminB1, HMGB1, p21) via immunocytochemistry (ICC) (Figure 1B). Then, an automated microscope (Eclipse Ti-PFS, Nikon) was employed to acquire thousands of images, which were further processed using the previously established method (ref) to segmented individual cells in the images and extract their intensity profiles of the stained senescence markers. This automated imaging and analysis process allow us to effectively profile tens of thousands of cells in the experimenting conditions (Figure 1C).

**Figure 1.**
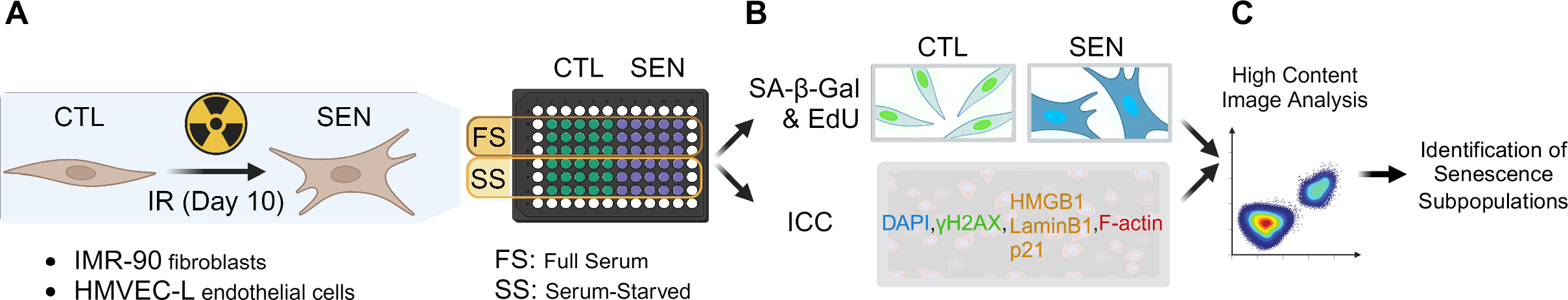
**High-content imaging workflow**. **a**) Sample preparation. Human lung primary microvascular endothelial cells (HMVEC-L) and fibroblasts (IMR-90) were induced to senescence using ionizing radiation (IR). IR and mock-irradiated cells (CTL) were either cultured in full-serum medium the entire time (FS) or switched to low-serum medium for the last 3 days of culture to induce quiescence in CTL cells (SS). **b**) Staining for senescence markers. Prepared samples were either co-stained for senescence-associated beta-galactosidase activity (SA-β-Gal) and proliferation via EdU incorporation (EdU); or for other senescence markers (γH2AX, LaminB1, HMGB1, p21) using immunocytochemistry (ICC). **c**) High-content image analysis was performed to identify senescent subpopulations.

First, this dataset was used to assess senescence-associated markers at the cell population level (Figure 2). As expected, in both investigated cell types, the SA-β-Gal staining was higher in IR cells compared to their respective CTL cells and EdU incorporation was lower (Figure 2B). Importantly, the EdU signal was low in CTL samples in SS conditions (as were senescent IR cells), indicating the SS CTL cells were indeed quiescent. When comparing senescent cells (IR) vs control, an increased number of nuclear γH2AX (gamma- phosphorylated H2A Histone Family, Member X) foci and higher p21 nuclear levels were observed using ICC staining, while both LaminB1 and HMGB1 nuclear staining were lower in senescent cells. This was true both when comparing IR-induced senescent cells to proliferating CTL cells (FS conditions) and IR cells to quiescent CTL cells (SS conditions). Taken together, these data confirm the successful senescence induction in IR samples. Additionally, differences in marker expression were observed between irradiated HMVEC-L endothelial cells and IMR-90 fibroblasts, i.e., SA-β-Gal levels were about 5-fold higher in senescent HMVEC-L compared to senescent IMR-90, while the p21 levels were lower in senescent HMVEC-L compared to senescent IMR-90. This observation already demonstrated the existence of heterogeneity between senescent cell populations that originate from different cell types.

**Figure 2.**
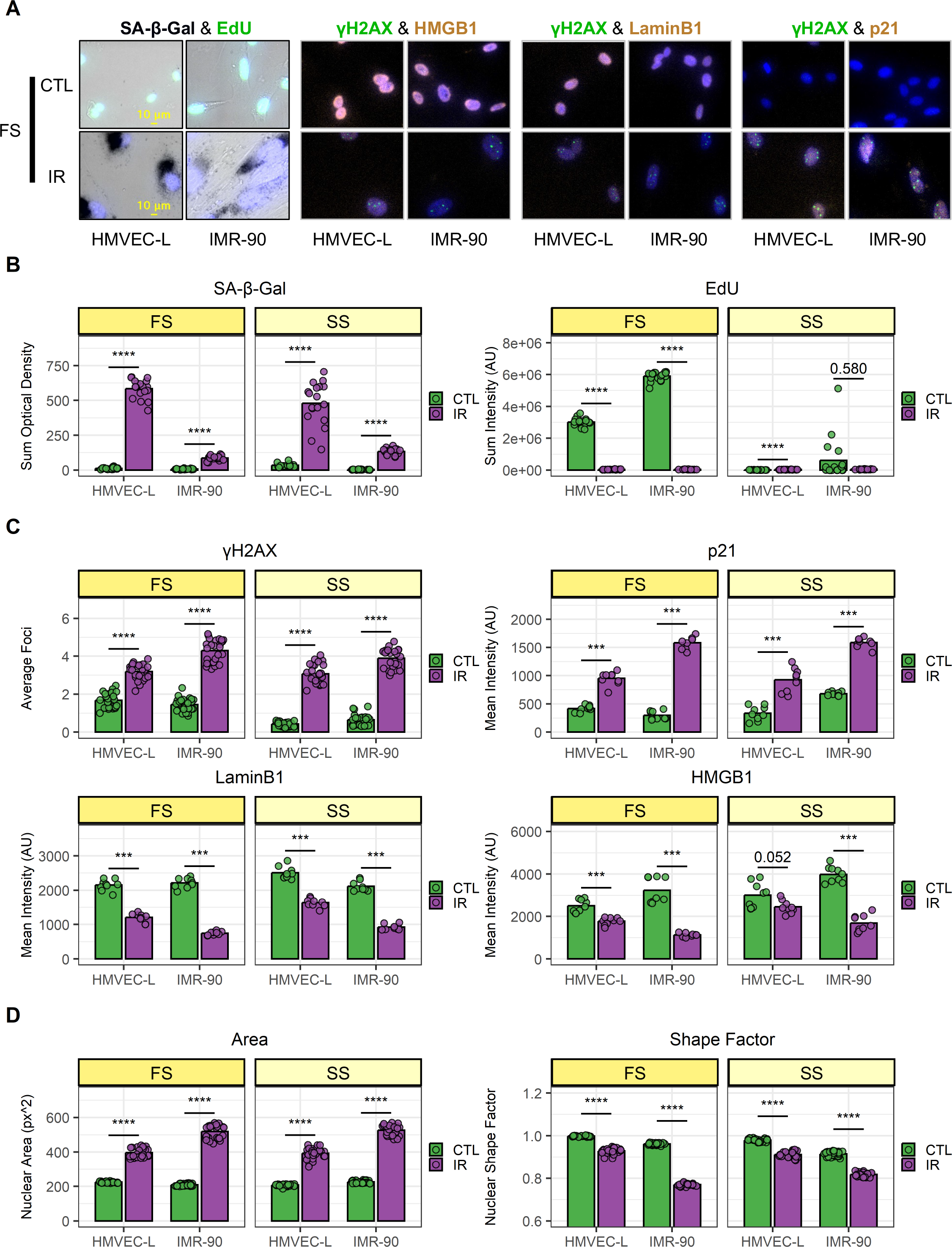
Validation of senescence induction and population-level heterogeneity**. a**) Representative images of senescence marker staining from full-serum (FS) samples. Top: mock-irradiated cells (CTL); bottom: ionizing-radiation-induced senescent cells (IR). For each co-staining, endothelial cells (HMVEC-L) are shown on the left; fibroblasts (IMR-90) are shown on the right. **b-d**) Image quantification of HMVEC-L and IMR-90 cells for both FS and serum- starved (SS) conditions. CTL samples are in green, while IR samples are in purple. Data shown are from 2 independent experiments. **b**) SA-β-Gal (left) and EdU (right) staining quantification. Each data point corresponds to one well (n = 18); bars indicate mean values. **c**) Immunocytochemistry staining quantification. Top-left: γH2AX; top-right: p21; bottom-left: LaminB1; bottom-right: HMGB1. Each data point corresponds to one well (γH2AX n = 27; p21, LaminB1, and HMGB1 n = 9); bars indicate mean values. **d**) Nuclear morphology feature quantification. Left: nuclear area; right: shape factor. Each data point corresponds to one well (n = 27); bars indicate mean values. ***: p-value < 10^-3^ ; ****: p-value < 10^-4^; non-significant values (p-value > 0.05) are shown.

### G2-arrested senescent cells express higher levels of senescence markers than G1- arrested senescent cells

After comparing population-level data, an exploratory analysis was performed within senescent cell populations using single-cell measurements of the markers assayed (Figure 3). Almost all of the co-staining experiments revealed two subpopulations with distinct senescence marker expression in IR-induced senescent endothelial cells (Figure 3A). It has been shown that the cell cycle is associated with cell phenotypes and protein expression^20^ ^21^. We hypothesized that these differences might be related to the phase of the cell cycle at which senescent cells were growth-arrested. It has been shown in our previous studies that DNA content and cell cycle phases can be identified from the intensity of fluorescently labeled nuclei in images^21–23^.

**Figure 3.**
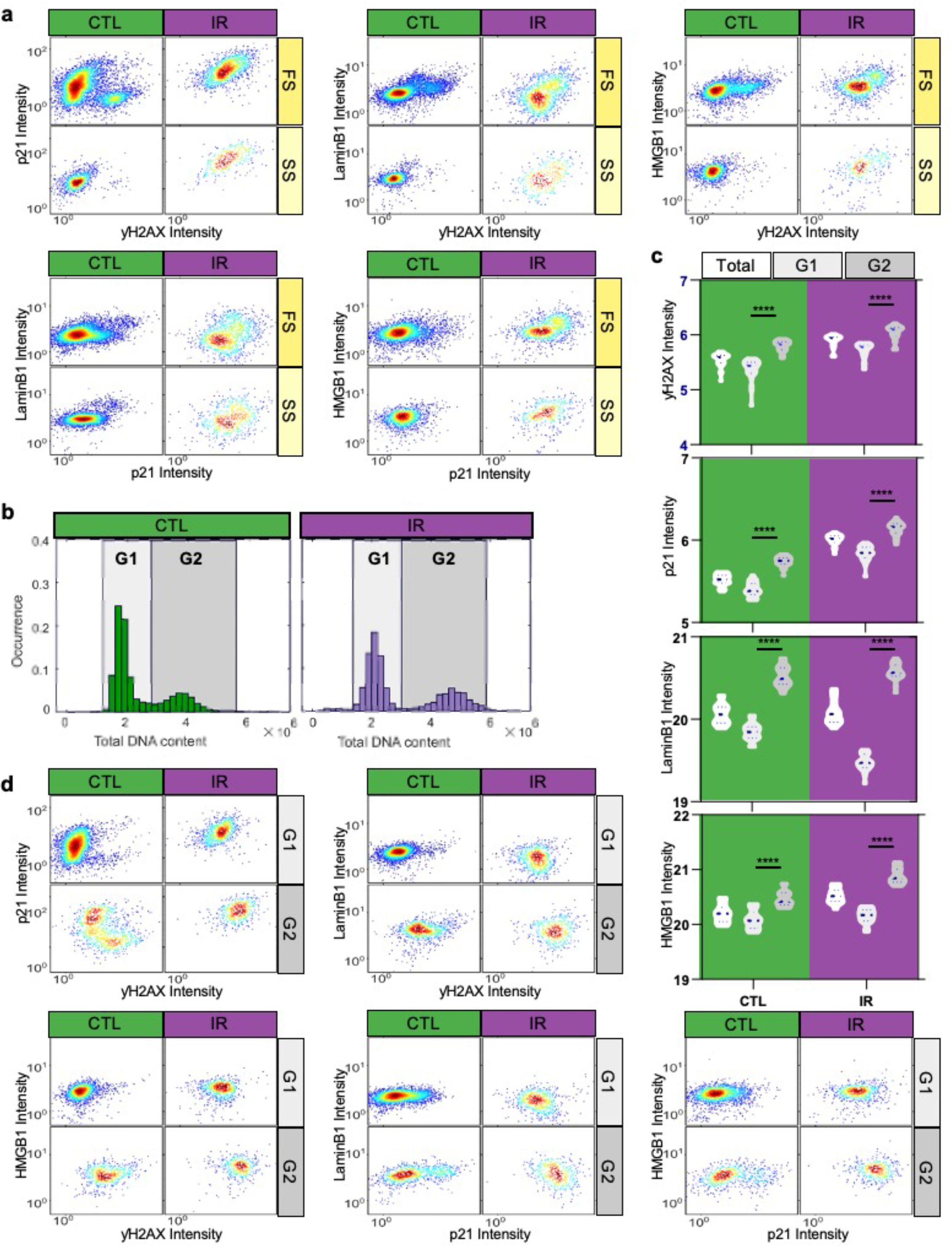
**Differential Expression of Senescence Markers in G1 and G2 Senescent Cells. a**) Density scatter plots display the expression levels of co-stained senescence markers in individual cells identified through immunofluorescence. The markers include P21 vs. γH2AX, Lamin B1 vs. γH2AX, HMGB1 vs. γH2AX, Lamin B1 vs. P21, and HMGB1 vs. P21 at the single-cell level under both IR and control conditions. Colors indicate density regional distribution density within the scatter plot. **b**) Histograms illustrate the distribution of total DNA content in individual cells under control and IR-treated conditions. DNA content is measured by the DAPI staining intensity in cell nuclei. The G1 and G2 cell cycle states are differentiated based on DAPI staining intensity. **c**) Violin plots present the average expression levels of various senescence markers from replicated wells and experiments within the overall population, as well as specifically in G1 or G2 states. A one-way ANOVA statistical test was performed where *: p-value < 0.05; **: p-value < 0.01; ***: p-value < 0.001; ****: p-value < 0.0001. The expression levels of senescence markers significantly differ between cell cycle states. **d**) Density scatter plots show the single-cell expression levels of various pairs of senescence markers in cells at G1 and G2 states, respectively.

To test this hypothesis, senescence marker staining (γH2AX, p21, LaminB1, and HMGB1) was analyzed in relation to DNA content (measured via DAPI staining), separating senescent endothelial cells into G1- and G2-arrested subpopulations based on low vs high DNA content (Figure 3B). Indeed, G1 and G2 senescent cells expressed different senescence marker levels, with G2-arrested senescent cells showing higher marker staining compared to G1-arrested cells (Figure 3C). Additionally, plotting G1 and G2 senescent cells separately resulted in uniformly stained subpopulations (Figure 3D), suggesting DNA content could be the main driver of the observed heterogeneity at the population level. Supplemental Figure 1 and Supplemental Figure 2 show similar observations for IMR-90 fibroblast cells.

### G1- and G2-arrested senescent endothelial cells show different IL-6 secretion and ABT263 susceptibility

Based on the differences in senescence marker expression, we hypothesized that G1- and G2-arrested senescent cells might also be functionally distinct. To test this hypothesis, we focused on senescent endothelial cells, where differences between G1- and G2-arrested cells were more prominent.

First, we optimized procedures to enrich senescent endothelial cell populations in G1 or G2 (Figure 4) according to previous reports^24^. We hypothesized that enriching cells in G2 or G1 just before irradiation would result in senescent populations enriched in G2 or G1 respectively (Figure 4A). Thus, G2- and G1-enriched populations were generated by either precisely timing cell seeding to obtain cells in their exponential growth phase (IR-G2-E) or using serum-starved culturing conditions (IR-G1-E), respectively. To compare the DNA content of senescent cells with that of cells at the time of irradiation (just before senescence induction), additional samples were prepared in parallel, which were fixed instead of being irradiated (CTL-G2-E, CTL-G1-E). As expected, the G2-E samples showed a higher percentage of G2 cells than the G1-E samples (Figure 4B). This was the case both at the time of irradiation (CTL samples, ∼40% vs 20% G2 cells) and, more importantly, in also in the senescent cell populations (IR samples, ∼60% vs 30% G2 cells). Interestingly, the relative difference in the percentage of G2 cells between G2-E and G1-E samples was about half for both CTL and IR samples (Figure 4C).

**Figure 4.**
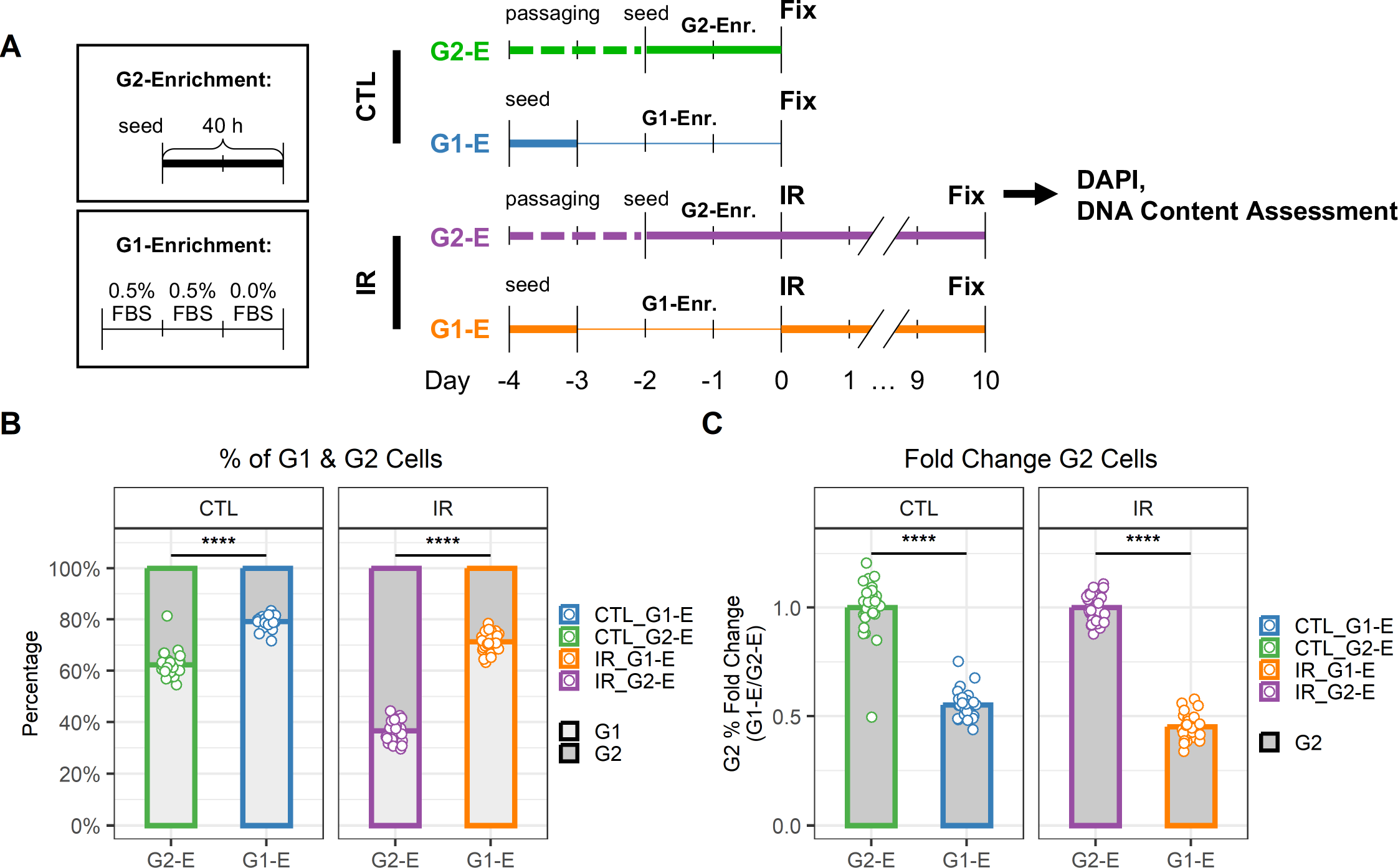
**G1 and G2 enrichment protocol for senescent endothelial cells**. **a**) Workflow to compare DNA content in cells just before irradiation (CTL) and senescent cells 10 days after irradiation (IR) when enriched for either G1 (G1-E) or G2 (G2-E) cells. **b**) Percentage of G1 and G2 cells per well. Each data point is a well (n = 30); bars indicate mean values. **c**) Fold change of G1 (left) and G2 (right) percentages in IR-G1-E vs IR-G2-E groups. Each data point is a well (n = 30); bars indicate mean values.

Upon establishing that DNA content enrichment at the time of senescence induction was maintained after cells became fully senescent, we proceeded to compare G2- and G1- enriched senescent populations (Figure 5). First, their IL-6 secretion levels were compared (Figure 5A). For this purpose, IR-G2-E and IR-G1-E samples were generated. Conditioned medium (CM) was collected over the last two days of culture, and then the samples were fixed and imaged. Using the imaging data, we validated the expected DNA content enrichment (Figure 5B-C). IL-6 secretion in the CM was measured by ELISA and normalized to cell counts obtained using imaging. IL-6 levels were higher in IR-G2-E compared to IR-G1-E samples (Figure 5D). Then, the sensitivity to ABT263 senolytic treatment was compared in IR-G2-E and IR-G1-E by measuring cell counts obtained via imaging (Figure 5E). DNA content enrichment was validated (Figure 5F), and viability was compared across different ABT263 concentrations (Figure 5G). Differences in ABT263 sensitivity were observed between IR-G2- E and IR-G1-E populations in two out of the three concentrations tested (0.33 and 1.00 µM). The same data were further analyzed to measure differences in viability within IR-G2-E and IR- G1-E populations by comparing their G1 and G2 subpopulations (Figure 5H). Differences in ABT263 sensitivity were observed between G1 and G2 subpopulations at all three concentrations tested (including 0.11 µM) both in the IR-G2-E and IR-G1-E populations. Thus, G2-arrested senescent endothelial cells secreted higher levels of IL-6 and were more sensitive to ABT263 senolytic treatment than G1-arrested cells. This suggests the existence of functionally distinct senescent cell subpopulations, which underscores the importance of considering senescence heterogeneity during the development of senotherapeutics.

**Figure 5.**
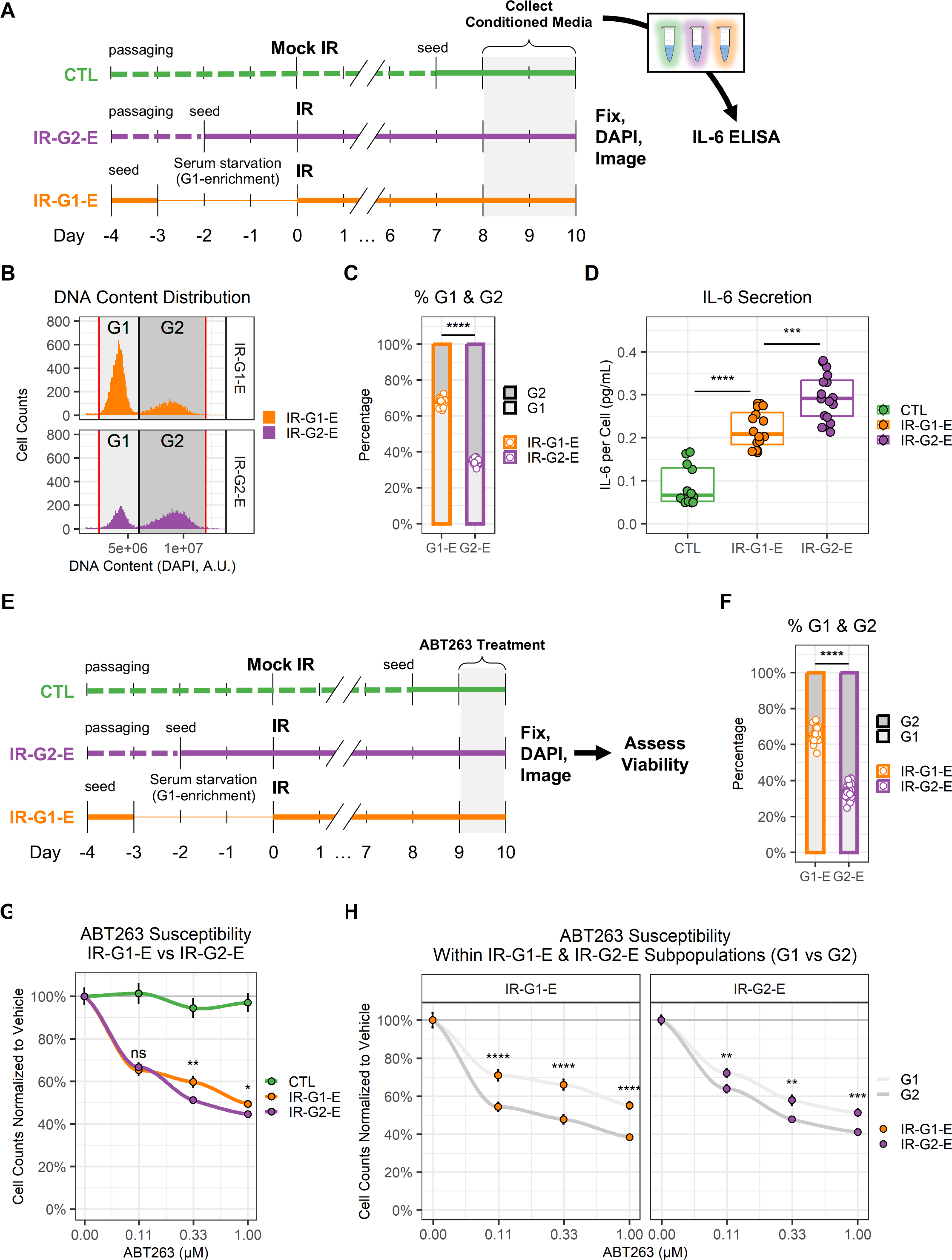
**G1 and G2 senescent endothelial cells show different levels of IL-6 secretion and ABT263 susceptibility**. **a**) Workflow for the comparison of IL-6 secretion in G1 vs G2 senescent endothelial cells. Three conditions were prepared: non-senescent mock- irradiated cells (CTL, green); ionizing radiation-induced senescent cells enriched in G2 (IR-G2- E, purple); ionizing radiation-induced senescent cells enriched in G1 (IR-G1-E, orange). Condition media (CM) were collected from the last 2 days of culture, after which cells were fixed, counterstained with DAPI, and imaged. IL-6 concentration was quantified by ELISA and normalized to cell counts. **b**) DNA content distribution histogram, showing G1 (light grey) and G2 (dark grey) peaks in IR-G2-E and IR-G1-E senescent populations. The plot shows all IR- G2-E and IR-G1-E cells from a single representative experiment. **c**) Quantification of (b), showing the sample percentages of senescent cells arrested in G1 and G2. Data shown is from 3 independent experiments; each data point is a sample (CTL n = 12; IR-G1-E and IR- G2-E n = 16). **d**) IL-6 secretion across CTL, IR-G2-E, and IR-G1-E groups normalized to cell counts. Data shown are from 3 independent experiments; each data point is a sample (CTL n = 12; IR-G1-E and IR-G2-E n = 16). ***: p-value < 10^-3^; ***: p-value < 10^-4^; by one-way ANOVA followed by post-hoc Tukey’s test. **e**) Workflow for the comparison of ABT263 susceptibility in G1 vs G2 senescent endothelial cells. CTL, IR-G2-E, and IR-G1-E were prepared as described in (a), but cells were treated with ABT263 for the last 24 h of culture. After treatment, cells were fixed, counterstained with DAPI, and imaged. **f**) Percentages of senescent cells per well arrested in G1 and G2. Data shown are from 3 independent experiments; each data point is a well (n = 30). **g**) Cell viability comparison after ABT263 treatment between IR-G2-E and IR-G1- E senescent populations, measured by cell counts normalized to vehicle (0.00 µM ABT263). Data shown are mean ± SEM for each ABT263 concentration from 3 independent experiments. Viability was compared between IR-G2-E and IR-G1-E populations across all ABT263 concentrations (n = 30). ns: p-value > 0.05; *: p-value < 0.05; **: p-value < 0.01; by non- parametric Mann-Whitney test corrected for multiple comparisons by FDR method. **h**) Cell viability comparison of G1 and G2 subpopulations within IR-G2-E and IR-G1-E populations from (g). Data shown are mean ± SEM of G1 (light grey) and G2 (dark grey) subpopulations for each ABT263 concentration from 3 independent experiments. Viability was compared between G1 and G2 cells across all ABT263 concentrations (n = 30). **: p-value < 10^-2^; ***: p-value < 10^-3^; ****: p-value < 10^-4^; by non-parametric Mann-Whitney test corrected for multiple comparisons by FDR method.

## Discussion

In this study, we aimed to identify functionally distinct subpopulations of senescent cells by using high-content imaging workflows. Specifically, our goal was to establish whether we could identify subpopulations of senescent cells with heterogeneous sensitivity to senolytic treatment.

By leveraging cell culture senescence models and analyzing single-cell measurements of several senescence-associated markers, we found a relationship between marker expression and DNA content. Specifically, we observed that G2-arrested senescent cells had higher levels of senescence markers than G1-arrested cells. Additionally, we found that G2- arrested senescent cells secreted higher levels of IL-6 and were more sensitive to ABT263 senolytic treatment. This suggests that DNA content can differentiate senescent subpopulations with distinct functions, as highlighted by senescence marker expression, SASP factor secretion, and drug response. While our study focused on ABT263, we speculate that the cytotoxic effect of other senolytic compounds might be heterogeneous and depend on the DNA content of the treated cells. Interestingly, previous studies in cancer cells have highlighted that the cytotoxic profile of several drugs is influenced by the DNA content and cell cycle phase at the time of treatment, with some drugs preferentially targeting cells in G1 and others in G2^25^.

It is important to note that this study focused on the analysis of two cell culture models (primary human endothelial cells and fibroblasts) of DNA-damage-induced senescence. Future studies will investigate different cell types and different senescence inducers and – most importantly – we will investigate *in vivo* heterogeneity of senescent cell subpopulations and their response to senolytics. While our study focused on senolytics, other types of senotherapeutics are being developed, such as senomorphics. Senomorphics can alleviate senescence-related tissue dysfunction by attenuating the SASP, rather than eliminating senescent cells^5^. It would be interesting to investigate (in future studies) whether senomorphics also display different effectiveness in targeting heterogeneous senescent subpopulations.

In summary, we demonstrated the existence of functionally distinct senescent subpopulations in culture, which can be differentiated based on G1 and G2 DNA content (Figure 6). To the best of our knowledge, our data also constitute the first evidence of heterogenous senolytic response between subpopulations of senescent cells. These findings highlight the importance of studying senescent cell heterogeneity and that their diversity should be considered when developing senolytic treatments.

**Figure 6.**
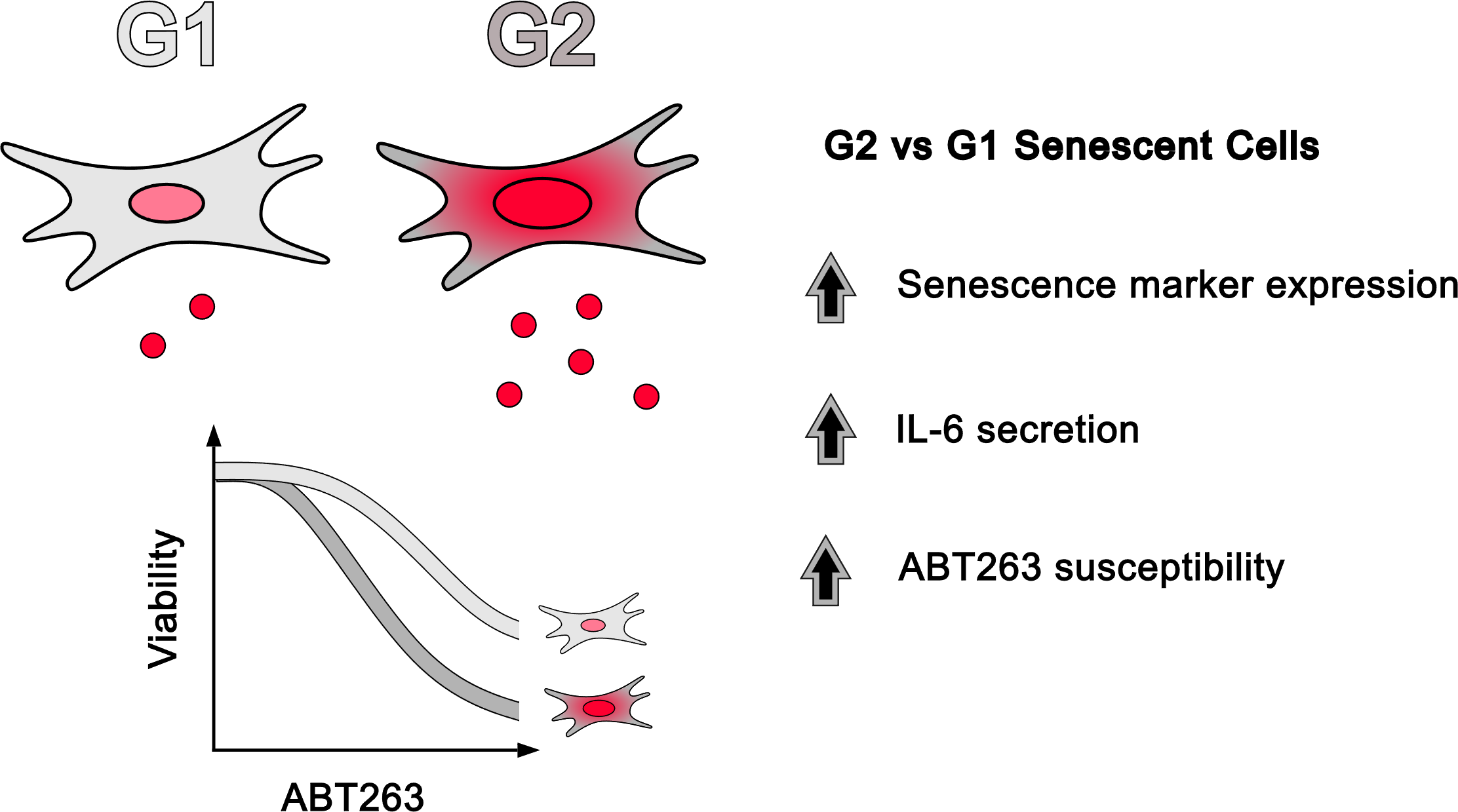
**Senescent heterogeneity model based on DNA content**. Compared to G1- arrested cells, G2-arrested senescent cells express higher levels of senescence-associated markers, secrete more IL-6, and are more sensitive to senolytic treatments.

## Materials and Methods

### Cell Culture

Primary human lung microvascular endothelial cells (HMVEC-L) were purchased from Lonza (CC-2527). HMVEC-L were cultured in EGM^TM^-2MV Microvascular Endothelial Cell Growth Medium-2 BulletKit^TM^ (Lonza, CC-3202) at 37°C, 14% O_2_, 5% CO_2_. Human lung fibroblasts IMR-90 were purchased from Coriell Institute (I90). IMR-90 cells were cultured in DMEM (Corning, 01-017-CV) supplemented with 10% FBS (R&D Systems, S11550H), 100 units/mL penicillin, and 100 µg/mL streptomycin (R&D Systems, B21210) at 37°C, 3% O_2_, 10% CO_2_. For all experiments, both HMVEC-L and IMR-90 were cultured in 96-well microplates appropriate for microscopy imaging (Corning, 3904), with media changes every 2-3 days. To achieve serum starvation in HMVEC-L, cells were washed twice in DPBS containing Ca^2+^ and Mg^2+^ (Gibco, 14040-117) and then cultured for 72 h in low-serum EGM^TM^-2MV medium (0.5% FBS instead of 5% FBS). To achieve serum starvation in IMR-90, cells were washed twice in DPBS containing Ca^2+^ and Mg^2+^ (Gibco, 14040-117) and then cultured for 72 h in low-serum DMEM medium (0.2% FBS instead of 10% FBS). To achieve G1-enrichment in HMVEC-L, cells were washed twice in DPBS containing Ca^2+^ and Mg^2+^ (Gibco, 14040-117) and then cultured for 48 h in low-serum EGM^TM^-2MV medium (0.5% FBS instead of 5% FBS) followed by 24 h in serum-free EGM^TM^-2MV medium. To achieve G2 enrichment in HMVEC-L, cells were seeded 40 h before irradiation was carried out.

### Senescence Induction

Senescence was induced as previously described by Neri et al.^24^. Briefly, cells were irradiated with 15 Gy, and medium change was performed immediately after treatment. Cells were considered senescent after at least 7 days since irradiation, during which medium was regularly changed (every 2-3 days).

### SA-**β**-Gal and EdU Staining

SA-β-Gal and EdU staining was performed following the Fully-Automated Senescence Test (FAST) workflow^26^. Briefly, commercially available kits were used to perform SA-β-Gal (Abcam, ab65351) and EdU staining (ThermoFisher Scientific, C10351). For EdU, cells were given medium containing 2.5 µM EdU 24 h before fixation. After 24 h, cells were fixed by adding 8% PFA in PBS pre-warmed to 37°C directly to the medium up to a final concentration of 4% PFA and incubated for 15 min at RT. Subsequently, cells were washed twice with PBS, and SA-β-Gal staining was performed.

To stain for SA-β-Gal, fixed cells were treated with the staining solution mix as recommended by the manufacturer (Abcam, ab65351). However, the X-Gal powder used was separately purchased (Life Technologies, 15520-018). Staining was performed overnight at 37°C in an incubator at atmospheric CO_2_ conditions. To prevent nonspecific indole crystal formation, empty spaces in between wells of the microplates were filled with PBS, and parafilm was used to seal the microplates before the overnight incubation. After the overnight incubation, cells were washed twice with PBS to stop the staining.

After SA-β-Gal staining, EdU detection was performed. Briefly, cells were permeabilized at room temperature for 15 min with 0.5% Triton X-100 (Millipore Sigma, T9284- 500ML) in PBS. After permeabilization, the Click-iT Reaction Cocktail was added as per the user manual, and cells were incubated for 30 min in the dark. After the incubation, cells were washed once with PBS, counterstained with 0.5 μg/mL DAPI in MilliQ water for 30 min at room temperature in the dark, then washed once with MilliQ water. Finally, wells were covered with PBS and imaged.

### Immunocytochemistry Staining

Immunocytochemistry staining was performed using standard protocols. Briefly, cells were first fixed and permeabilized as described in the SA-β-Gal and EdU staining section above. Then, cells were incubated with 10% goat serum for 1 h for blocking, incubated with primary antibodies overnight at 4°C, washed 3 times with PBS, and incubated with secondary antibodies for 1 h at room temperature in the dark. Afterward, samples were washed once with PBS, counterstained with 0.5 μg/mL DAPI in MilliQ water for 30 min at room temperature in the dark, then washed once with MilliQ water. Finally, wells were covered with PBS and imaged.

### ELISA

Conditioned medium (CM) of senescent and control cells was collected after 48 h culturing in fresh medium. Each sample for downstream ELISA analysis was obtained by pooling CM from at least three 96-well microplate wells of the same group (CTL, IR-G1-E, or IR-G2-E). To remove potential cell debris, CM was then centrifuged at 10,000 g for 1 min and the supernatant was moved to a new tube. A human IL-6 ELISA kit (Thermo Fisher Scientific, KHC0061) was then used as instructed by the manufacturer to measure IL-6 concentrations. To normalize IL-6 secretion, cells were fixed after collection of CM in 4 % PFA for 15 min at room temperature, washed twice with PBS, permeabilized with 0.5% Triton X-100 at room temperature for 15 min, washed twice with PBS, counterstained with 0.5 μg/mL DAPI in MilliQ water for 30 min at room temperature in the dark, then washed once with MilliQ water. Finally, wells were covered with PBS and imaged. DAPI staining was used to obtain cell counts per well, which were then used to normalize IL-6 concentrations of all samples.

### Senolytic Treatment

On the last day before analysis, cells were treated with the senolytic ABT263/Navitoclax (Selleck Chemicals, S1001) at different concentrations for 24 h, while only vehicle (DMSO) was given as mock treatment.

### Image Acquisition

Wide-field microscopy was performed on a Nikon Eclipse Ti-PFS fully motorized microscope controlled by NIS Elements AR 5.21 (Nikon, Melville, NY). The setup comprised a Lambda 10-3 emission filter wheel, a SmartShutter in the brightfield light path, and a 7-channel Lambda 821 (Sutter Instruments, Novato, CA) LED epifluorescence light source with excitation filters on the LEDs, controlled by a PXI 6723 DAQ (NIDAQ; National Instruments) board. Images were acquired by an Andor iXon Life 888 EMCCD camera (Oxford Instruments) using 10-100 ms exposure times, with a Nikon S Fluor 20× DIC NA=0.5 lens. To image SA-β-Gal staining, an incandescent Koehler illumination was used, and a 692/40 “emission” filter. The Kohler condenser was carefully focused for each experiment in the center of a well, with the aperture diaphragm semi-open. 5×5 tiled images were recorded without overlap or registration, using the full 1024×1024 resolution of the camera (1.3 µm/pixel). For autofocusing, the Nikon’s Perfect Focus System was used. Data were saved as native *.nd2 files for SA-β-Gal and EdU staining, or as *.tiff files for immunocytochemistry staining.

### Image Processing

For SA-β-Gal and EdU images, native format *.nd2 image files were opened in Image Analyst MKII (Image Analyst Software, Novato, CA). Analysis was performed using modified standard and custom pipelines described in the FAST workflow^26^. Fluorescence background was defined as the 20th percentile of the image intensity histogram. The output Excel file containing single-cell measurements for each whole microplate was saved, and further data analysis was performed in R.

For immunofluorescence images, a previously developed pipeline was used to analyze the data^18,23^. In short, images from individual fields of view (FOV) within the same well were first stitched together to form a single large FOV. DAPI and Phalloidin stained fluorescent channels were then used to obtain masks of individual nuclei and cells in the images using a custom segmentation pipeline^23^. These cell and nuclei masks were subsequently applied to all fluorescence channels with different molecular staining to extract the stained intensity profiles at the individual cell level. We also obtain the morphology features such as area, aspect ratio, shape factors for each segmented cell and nuclei masks^27^.

### Data Analysis

Data for Figures 2, 4, and 5 were analyzed with custom R pipelines, available on GitHub (https://github.com/f-neri/Wirtz-collaboration). For SA-β-Gal and EdU data, raw single-cell measurements were first pre-processed using an R Shiny-based application, FAST.R^26^. Suppl. Table S1, Suppl. Table S2, Suppl. Table S3, Suppl. Table S4, Suppl. Table S5, Suppl. Table S6, Suppl. Table S7, Suppl. Table S8, and Suppl. Table S9 show underlying data for figures.

### Statistics

Statistical tests employed are exhaustively described in each figure endothelial. Such statistical tests were either performed in R (v4.3.2) or MATLAB (R2023a).

## Abbreviations

PFA: paraformaldehyde
EdU: 5-ethynyl-2’-deoxyuridine
SA-β-Gal: senescence-associated β galactosidase activity
HMVEC-L: human microvascular endothelial cells from lung
CTL: control non-senescent cells
IR: ionizing-radiation induced senescent cells
IR-G1-E: G1-enriched IR cells
IR-G2-E: G2-enriched IR cells
FS: full serum
SS: serum starved
CM: conditioned medium

## Author Contributions

B.S., P.-H.W., D.W., and J.C. conceptualized and designed the study, and acquired funding for this project. F.N. devised and performed the experiments. F.N. and S.Z. performed data analysis, generated figures, and wrote the initial manuscript draft. A.A.G. and P.-H.W. developed the high-content imaging workflow utilized. M.W., P.-Y.D., A.A.G., D.W., P.-H.W., and B.S. reviewed and edited the manuscript.

## Acknowledgments

We would like to thank Dr. Corey Webster for helpful discussions regarding visual displays.

---

Supplemental Figure 1. High Content Imaging for IMR 90 Cells.

Supplemental Figure 2. Differential Expression of Senescence Markers in G1 and G2. Senescent Cells in IMR-90 cells. Violin plots present the average expression levels of various senescence markers from replicated wells and experiments within the overall population, G1 or G2 states. A one-way ANOVA statistical test was performed where ****: p- value < 0.0001. The expression levels of senescence markers significantly differ between cell cycle states.

## References

1 Gorgoulis, V. et al. Cellular Senescence: Defining a Path Forward. Cell 179, 813–827, doi:10.1016/j.cell.2019.10.005 (2019).

2 Xu, M. et al. Senolytics Improve Physical Function and Increase Lifespan in Old Age. Nature medicine 24, 1246, doi:10.1038/S41591-018-0092-9 (2018).

3 Baker, D. J. et al. Naturally occurring p16Ink4a-positive cells shorten healthy lifespan. Nature 530, 184–189, doi:10.1038/nature16932 (2016).

4 Chang, J. et al. Clearance of senescent cells by ABT263 rejuvenates aged hematopoietic stem cells in mice. Nature Medicine 22, 78–83, doi:10.1038/nm.4010 (2016).

5 Chaib, S., Tchkonia, T. & Kirkland, J. L. Cellular senescence and senolytics: the path to the clinic. Nature Medicine 28, 1556–1568, doi:10.1038/s41591-022-01923-y (2022).

6 Di Micco, R., Krizhanovsky, V., Baker, D. & d’Adda di Fagagna, F. Cellular senescence in ageing: from mechanisms to therapeutic opportunities. Nature Reviews Molecular Cell Biology 2020 22:*2* 22, 75-95, doi:10.1038/s41580-020-00314-w (2020).

7 Tsuruda, P. et al. UBX1325, a small molecule inhibitor of Bcl-xL, attenuates vascular dysfunction in two animal models of retinopathy.

8 UNITY Biotechnology Announces Positive 48-Week Results from Phase 2 BEHOLD Study of UBX1325 in Patients with Diabetic Macular Edema.

9 Cohn, R. L., Gasek, N. S., Kuchel, G. A. & Xu, M. The heterogeneity of cellular senescence: insights at the single-cell level. Trends in Cell Biology 33, 9–17, doi:10.1016/j.tcb.2022.04.011 (2023).

10 Coppé, J.-P. et al. Senescence-Associated Secretory Phenotypes Reveal Cell-Nonautonomous Functions of Oncogenic RAS and the p53 Tumor Suppressor. PLoS Biology 6, e301, doi:10.1371/journal.pbio.0060301 (2008).

11 Basisty, N. et al. A proteomic atlas of senescence-associated secretomes for aging biomarker development. PLOS Biology 18, e3000599, doi:10.1371/journal.pbio.3000599 (2020).

12 Cells, S. et al. Unmasking Transcriptional Heterogeneity in Senescent Cells. Current Biology 27, 2652–2660.e2654, doi:10.1016/j.cub.2017.07.033 (2017).

13 Ciotlos, S., Wimer, L., Campisi, J. & Melov, S. (bioRxiv, 2023).

14 Fard, A. T. et al. Deconstructing replicative senescence heterogeneity of human mesenchymal stem cells at single cell resolution reveals therapeutically targetable senescent cell sub-populations. (Bioinformatics, 2022).

15 Tang, H. et al. Single senescent cell sequencing reveals heterogeneity in senescent cells induced by telomere erosion. Protein & Cell 10, 370–375, doi:10.1007/s13238-018-0591-y (2019).

16 Zhu, Y. et al. The Achilles’ heel of senescent cells: from transcriptome to senolytic drugs. Aging Cell 14, 644–658, doi:10.1111/ACEL.12344 (2015).

17 Way, G. P., Sailem, H., Shave, S., Kasprowicz, R. & Carragher, N. O. Evolution and impact of high content imaging. SLAS Discov 28, 292–305, doi:10.1016/j.slasd.2023.08.009 (2023).

18 Wu, P. H. et al. Evolution of cellular morpho-phenotypes in cancer metastasis. Sci Rep 5, 18437, doi:10.1038/srep18437 (2015).

19 Dimri, G. P. et al. A biomarker that identifies senescent human cells in culture and in aging skin in vivo. Proc Natl Acad Sci U S A 92, 9363–9367, doi:10.1073/pnas.92.20.9363 (1995).

20 Chen, W. C. et al. Functional interplay between the cell cycle and cell phenotypes. Integr Biol (Camb) 5, 523–534, doi:10.1039/c2ib20246h (2013).

21 Chambliss, A. B., Wu, P. H., Chen, W. C., Sun, S. X. & Wirtz, D. Simultaneously defining cell phenotypes, cell cycle, and chromatin modifications at single-cell resolution. FASEB J 27, 2667–2676, doi:10.1096/fj.12-227108 (2013).

22 Wu, P. H., Hung, S. H., Ren, T., Shih Ie, M. & Tseng, Y. Cell cycle-dependent alteration in NAC1 nuclear body dynamics and morphology. Phys Biol 8, 015005, doi:10.1088/1478-3975/8/1/015005 (2011).

23 Wu, P. H. et al. Single-cell morphology encodes metastatic potential. Sci Adv 6, eaaw6938, doi:10.1126/sciadv.aaw6938 (2020).

24 Neri, F., Basisty, N., Desprez, P. Y., Campisi, J. & Schilling, B. Quantitative Proteomic Analysis of the Senescence-Associated Secretory Phenotype by Data-Independent Acquisition. Current Protocols 1, e32, doi:10.1002/cpz1.32 (2021).

25 Johnson, T. I. et al. Quantifying cell cycle-dependent drug sensitivities in cancer using a high throughput synchronisation and screening approach. eBioMedicine 68, doi:10.1016/j.ebiom.2021.103396 (2021).

26 Neri, F. et al. A Fully-Automated Senescence Test (FAST) for the high-throughput quantification of senescence-associated markers. Geroscience, doi:10.1007/s11357-024-01167-3 (2024).

27 Phillip, J. M. et al. Biophysical and biomolecular determination of cellular age in humans. Nat Biomed Eng 1, doi:10.1038/s41551-017-0093 (2017).

