## Supplemental Figure 1 for "Senescent cell heterogeneity and responses to senolytic treatment are related to cell cycle status during cell growth arrest"

### Slide 1
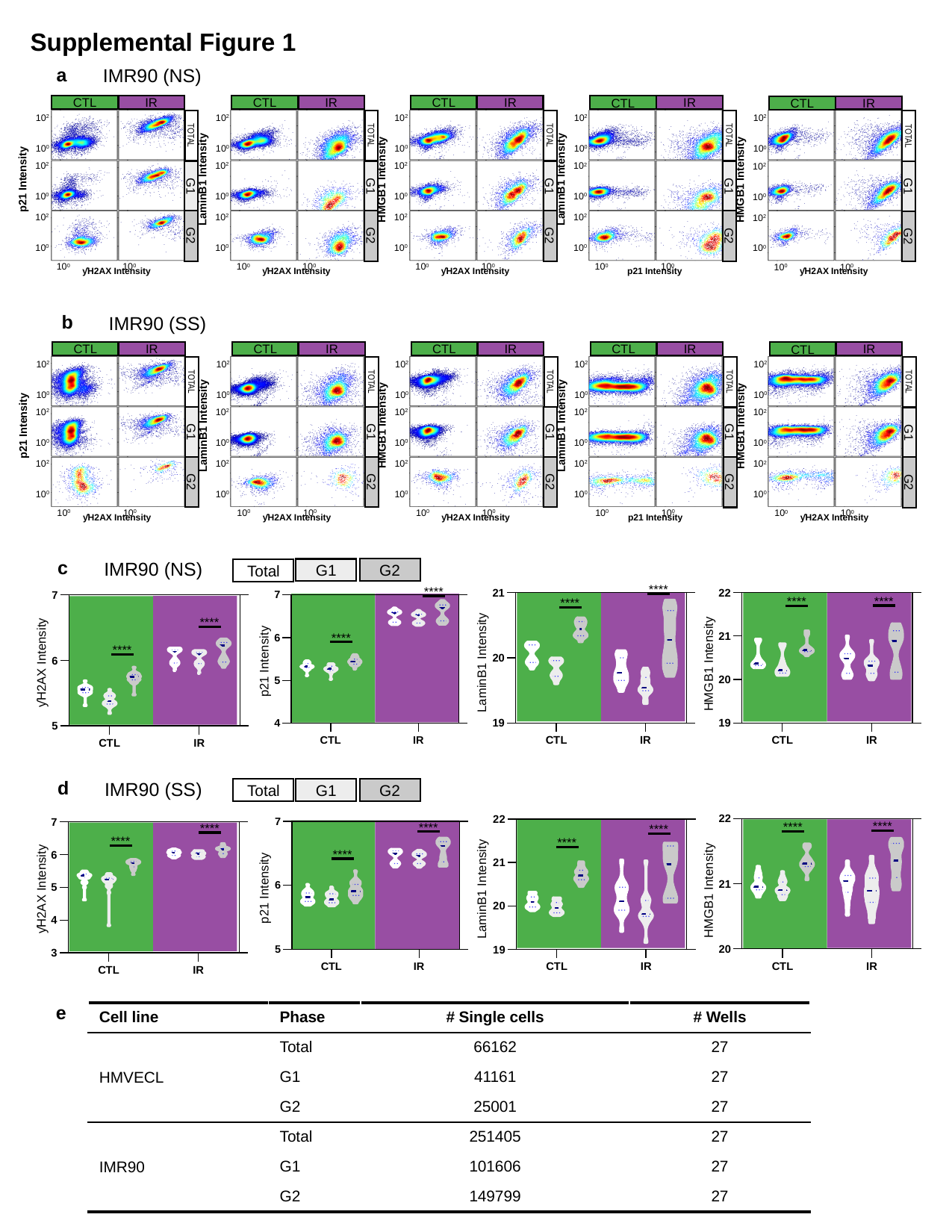

Supplemental Figure 1
a
IMR90 (NS)
IR
CTL
TOTAL
G1
G2
p21 Intensity
ƴH2AX Intensity
10^2^
10^0^
10^2^
10^0^
10^2^
10^0^
10^0^
10^0^
IR
CTL
TOTAL
G1
G2
LaminB1 Intensity
ƴH2AX Intensity
10^2^
10^0^
10^2^
10^0^
10^2^
10^0^
10^0^
10^0^
IR
CTL
TOTAL
G1
G2
HMGB1 Intensity
ƴH2AX Intensity
10^2^
10^0^
10^2^
10^0^
10^2^
10^0^
10^0^
10^0^
IR
CTL
TOTAL
G1
G2
LaminB1 Intensity
p21 Intensity
10^2^
10^0^
10^2^
10^0^
10^2^
10^0^
10^0^
10^0^
IR
CTL
TOTAL
G1
G2
HMGB1 Intensity
ƴH2AX Intensity
10^2^
10^0^
10^2^
10^0^
10^2^
10^0^
10^0^
10^0^
b
IMR90 (SS)
IR
CTL
TOTAL
G1
G2
p21 Intensity
ƴH2AX Intensity
10^2^
10^0^
10^2^
10^0^
10^2^
10^0^
10^0^
10^0^
IR
CTL
TOTAL
G1
G2
LaminB1 Intensity
ƴH2AX Intensity
10^2^
10^0^
10^2^
10^0^
10^2^
10^0^
10^0^
10^0^
IR
CTL
TOTAL
G1
G2
HMGB1 Intensity
ƴH2AX Intensity
10^2^
10^0^
10^2^
10^0^
10^2^
10^0^
10^0^
10^0^
IR
CTL
TOTAL
G1
G2
LaminB1 Intensity
p21 Intensity
10^2^
10^0^
10^2^
10^0^
10^2^
10^0^
10^0^
10^0^
IR
CTL
TOTAL
G1
G2
HMGB1 Intensity
ƴH2AX Intensity
10^2^
10^0^
10^2^
10^0^
10^2^
10^0^
10^0^
10^0^
c
IMR90 (NS)
G2
G1
Total
****
****
****
****
****
****
****
****
p21 Intensity
ƴH2AX Intensity
LaminB1 Intensity
HMGB1 Intensity
d
IMR90 (SS)
G2
G1
Total
****
****
****
****
****
****
****
****
p21 Intensity
ƴH2AX Intensity
LaminB1 Intensity
HMGB1 Intensity
e
| Cell line | Phase | # Single cells | # Wells |
| --- | --- | --- | --- |
| HMVECL | Total | 66162 | 27 |
| | G1 | 41161 | 27 |
| | G2 | 25001 | 27 |
| IMR90 | Total | 251405 | 27 |
| | G1 | 101606 | 27 |
| | G2 | 149799 | 27 |

### Slide 2
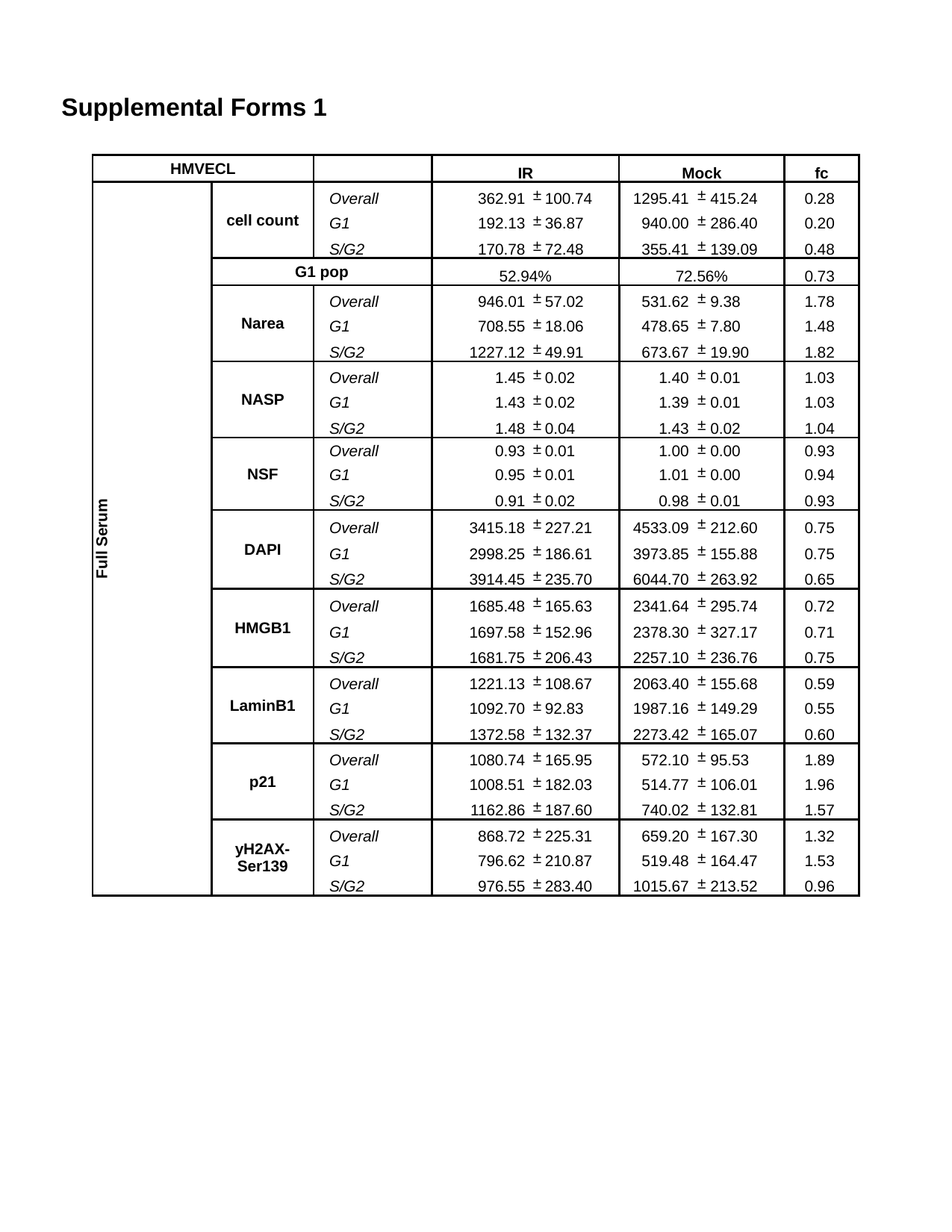

Supplemental Forms 1
| HMVECL | | | IR | | | Mock | | | fc |
| --- | --- | --- | --- | --- | --- | --- | --- | --- | --- |
| Full Serum | cell count | Overall | 362.91 | ± | 100.74 | 1295.41 | ± | 415.24 | 0.28 |
| | | G1 | 192.13 | ± | 36.87 | 940.00 | ± | 286.40 | 0.20 |
| | | S/G2 | 170.78 | ± | 72.48 | 355.41 | ± | 139.09 | 0.48 |
| | G1 pop | | 52.94% | | | 72.56% | | | 0.73 |
| | Narea | Overall | 946.01 | ± | 57.02 | 531.62 | ± | 9.38 | 1.78 |
| | | G1 | 708.55 | ± | 18.06 | 478.65 | ± | 7.80 | 1.48 |
| | | S/G2 | 1227.12 | ± | 49.91 | 673.67 | ± | 19.90 | 1.82 |
| | NASP | Overall | 1.45 | ± | 0.02 | 1.40 | ± | 0.01 | 1.03 |
| | | G1 | 1.43 | ± | 0.02 | 1.39 | ± | 0.01 | 1.03 |
| | | S/G2 | 1.48 | ± | 0.04 | 1.43 | ± | 0.02 | 1.04 |
| | NSF | Overall | 0.93 | ± | 0.01 | 1.00 | ± | 0.00 | 0.93 |
| | | G1 | 0.95 | ± | 0.01 | 1.01 | ± | 0.00 | 0.94 |
| | | S/G2 | 0.91 | ± | 0.02 | 0.98 | ± | 0.01 | 0.93 |
| | DAPI | Overall | 3415.18 | ± | 227.21 | 4533.09 | ± | 212.60 | 0.75 |
| | | G1 | 2998.25 | ± | 186.61 | 3973.85 | ± | 155.88 | 0.75 |
| | | S/G2 | 3914.45 | ± | 235.70 | 6044.70 | ± | 263.92 | 0.65 |
| | HMGB1 | Overall | 1685.48 | ± | 165.63 | 2341.64 | ± | 295.74 | 0.72 |
| | | G1 | 1697.58 | ± | 152.96 | 2378.30 | ± | 327.17 | 0.71 |
| | | S/G2 | 1681.75 | ± | 206.43 | 2257.10 | ± | 236.76 | 0.75 |
| | LaminB1 | Overall | 1221.13 | ± | 108.67 | 2063.40 | ± | 155.68 | 0.59 |
| | | G1 | 1092.70 | ± | 92.83 | 1987.16 | ± | 149.29 | 0.55 |
| | | S/G2 | 1372.58 | ± | 132.37 | 2273.42 | ± | 165.07 | 0.60 |
| | p21 | Overall | 1080.74 | ± | 165.95 | 572.10 | ± | 95.53 | 1.89 |
| | | G1 | 1008.51 | ± | 182.03 | 514.77 | ± | 106.01 | 1.96 |
| | | S/G2 | 1162.86 | ± | 187.60 | 740.02 | ± | 132.81 | 1.57 |
| | yH2AX-Ser139 | Overall | 868.72 | ± | 225.31 | 659.20 | ± | 167.30 | 1.32 |
| | | G1 | 796.62 | ± | 210.87 | 519.48 | ± | 164.47 | 1.53 |
| | | S/G2 | 976.55 | ± | 283.40 | 1015.67 | ± | 213.52 | 0.96 |
