## Supplemental Figure 2 for "Senescent cell heterogeneity and responses to senolytic treatment are related to cell cycle status during cell growth arrest"

### Slide 1
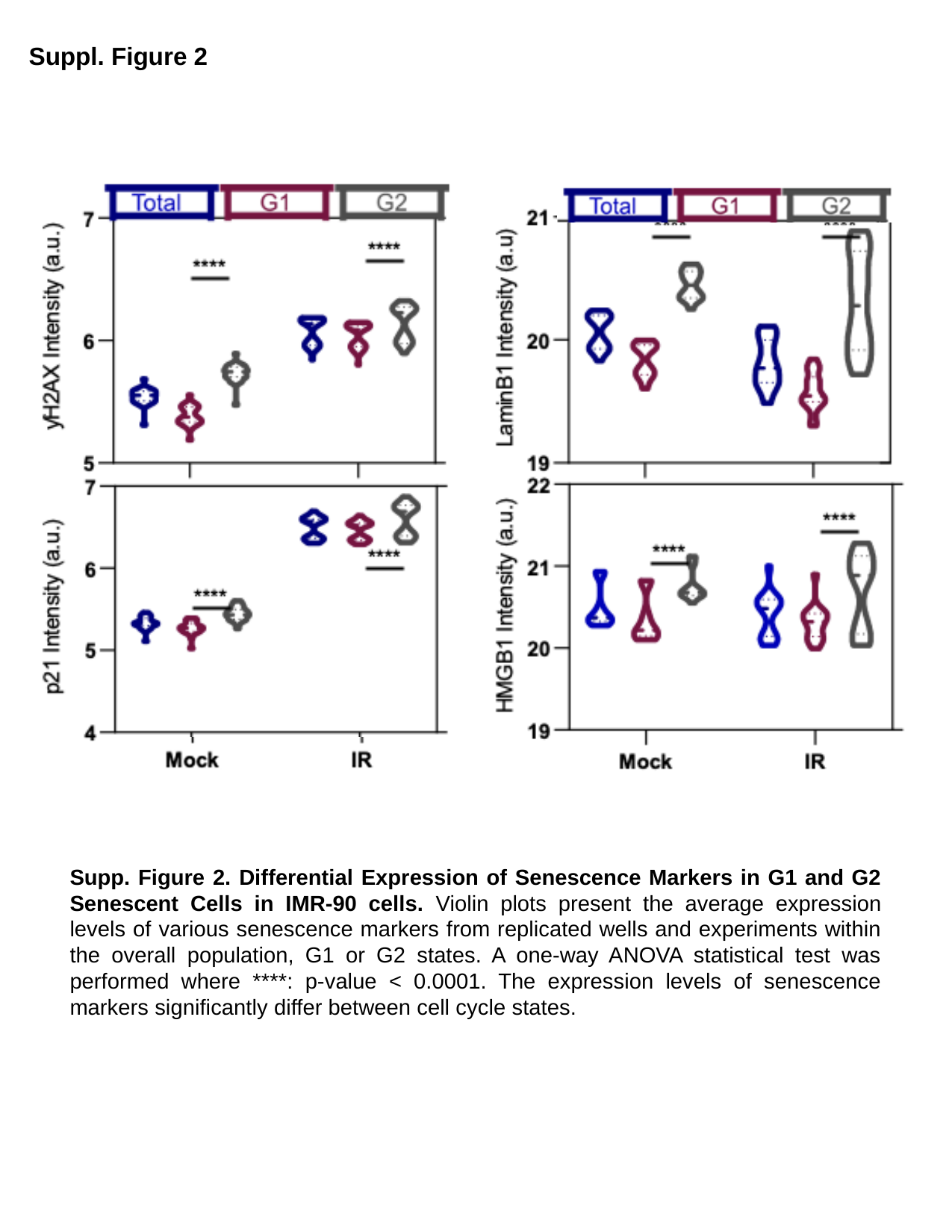

Suppl. Figure 2
Supp. Figure 2. Differential Expression of Senescence Markers in G1 and G2 Senescent Cells in IMR-90 cells. Violin plots present the average expression levels of various senescence markers from replicated wells and experiments within the overall population, G1 or G2 states. A one-way ANOVA statistical test was performed where ****: p-value < 0.0001. The expression levels of senescence markers significantly differ between cell cycle states.
